# Variation of wine preference amongst consumers is influenced by the composition of salivary proteins

**DOI:** 10.1101/2023.03.27.534455

**Authors:** Jiaqiang Luo, Xinwei Ruan, Ching-Seng Ang, Yada Nolvachai, Philip J. Marriott, Pangzhen Zhang, Kate Howell

**Affiliations:** School of Agriculture and Food Sciences, Faculty of Science, The University of Melbourne, Parkville, Australia; Melbourne Mass Spectrometry and Proteomics Facility, Bio21 Molecular Science and Biotechnology Institute, The University of Melbourne, Parkville, Australia; Australian Centre for Research on Separation Science, School of Chemistry, Monash University, Clayton Australia

**Author notes:** Corresponding Author: Associate Professor Kate Howell.

**Keywords:** Shiraz, Wine flavour perception, Saliva, Volatiles, Salivary proteins

## Abstract

The preferences of consumers for different flavours and aromas in wine are varied and may be explained by inherent factors such as cultural background, wine education and personal taste of the wine consumer. Wine flavour as perceived in the mouth includes aroma compounds released through the retronasal pathway which are shaped by interactions with saliva. Saliva and wine interactions could provide an explanation as to why wine tasters express different preferences for wine. To test this hypothesis, 13 Western and 13 Chinese experienced wine tasters were recruited. Sensory evaluation was performed in formal surroundings to acquire free description-based and perceived sensory intensity data using the Pivot^®^ Profile and continuous scale assessment, respectively. Participants’ saliva samples were collected before the sensory evaluation and spiked into a wine sample to investigate the impact on the wine volatile release using GC×GC−MS. Saliva samples were subjected to enzyme activity assays and protein composition profiling by Tandem Mass Tag (TMT) quantitative proteomics. The wine tasters showed differences in wine flavour perception, which was supported by the difference in wine volatile release resulting from the addition of saliva. The two groups of participants did not have significant differences in total salivary protein concentrations or the amounts of esterase and α-amylase. However, statistically significant variations in the concentrations of specific proteins (proline-rich proteins (PRPs) and lipocalin-1 (LCN-1); *p* < 0.01) were found between the two groups. Significant correlations between perceived intensities of wine attributes and concentrations of PRPs and LCN-1 were observed. These results indicate that the composition of proteins in saliva are a factor that influences wine perception and preference. Our results provide a biochemical basis to understanding preference for food based on interactions between aroma compounds and salivary proteins and could be used to suggest foods or beverages to particular cultural groups.

## INTRODUCTION

Food preference can be shaped by gender, age, body weight and cultural background.^1^ Amongst these factors, culture is particularly influential, as it determines the native cuisine on which an individual is raised.^1,2^ Environment and agricultural practices are the principal cultural factors that impacts upon people’s food choices, as geographic and climatic conditions influence the availability of plants and animals.^2^ Other factors such as personal and religious beliefs, social and family status, levels of innovation, mechanisation and experimentation, and transportation are also influential.^2^ The variation of food preference can potentially be better understood with formal sensory evaluation combined with chemical analyses on compounds that contribute to sensory perception. However, there are more mechanical variables to consider when analysing perception and preference for solid food samples in a formal sensory analysis^3^ where controlling mastication and swallowing is difficult to achieve. Wine is a suitable food liquid to investigate the cultural preference for food in a controlled environment.

Stylistic preference for wine is influenced by a consumer’s cultural background, level of wine education and personal taste.^4^ Amongst sensations during wine tasting, the perception of wine aromas through orthonasal and retronasal olfaction is considered the most influential for the quality rating of wine.^5^ Aromatic compounds undergo complicated interactions in the retronasal pathway. These interactions can occur as soon as the wine enters the oral cavity and after swallowing.^6^ Saliva can alter the partitioning of wine volatiles by dilution,^7^ salting out,^8^ and modification of the rheological properties of the wine.^9^ On the other hand, salivary components are able to participate in multiple interactions. α-Amylase and mucin, the predominant proteins in human saliva, demonstrate a strong volatile binding capacity for esters by hydrophobic interactions.^10^ Salivary esterases,^10^ aldehyde dehydrogenases,^11^ and peroxidases^12^ can easily access their substrates in wine and hence modify the wine volatile profile in the mouth. As oral microorganisms contribute to the hydrolysis of glucosides in wine,^13^ there is a role for microbial contribution to aroma release during wine tasting.^14^

The composition of saliva determines how an individual perceives wine. Psychological stress,^15^ smoking,^16^ caffeine intake,^17^ exercise,^17^ and dietary patterns^18^ alter salivary parameters such as flow rate, total protein concentration, protein profile, and microbiota of resting and unstimulated saliva with the impact of diet have been observed to affect saliva. These factors may be influenced by ethnicity; Zhang et al.^19^ studied 304 Chinese participants from two ethnicities (200 Han and 104 Mongolian) revealing that the Mongolian group had significantly higher lipolytic activities in saliva samples while unstimulated salivary α-amylase activity was significantly lower. The authors explained the variations as associated with the distinct dietary habits between the two groups. The Mongolian diet is higher in fat intake while the Han diet is more abundant in carbohydrates. Mosca et al.^20^ reported that unstimulated saliva in Chinese donors (*n* = 15) had significantly higher salivary protein concentrations than Caucasian donors (*n* = 15). However, significant differences in other salivary parameters, including salivary flow rate, as well as amylolytic and lipolytic activities, were not observed, and a more comprehensive unbiased proteomics profiling analysis was not performed. Diet may be responsible for differences in saliva content as observed by Mennella et al.^21^ who investigated 42 Italian participants and observed a positive correlation between the high-fat diet and lipase activity. Rossi et al.^22^ demonstrated that Qataris of Arab descent (*n* = 1518) had significantly lower salivary amylase gene *AMY1* copy numbers than Qataris of Persian descent (*n* = 948), which could be associated with their difference in dietary starch intake.^23^ Previous studies by our group have proposed that the different prevalence of *Veillonella* in saliva samples of Western (*n* = 13) and Chinese (*n* = 13) wine tasters could be a possible contributor to their differential perception of bitterness and astringency^14^. The abundance of *Veillonella* in the saliva was positively correlated to the abundance of “flavone and flavonol biosynthesis” pathway and wine flavonols are associated with these two sensory attributes.^14^

Physiological differences amongst consumers with different cultural backgrounds are not likely to contribute to the variation in food preferences,^2,24^ and recent studies have demonstrated that saliva and aroma compounds interact in different groups of individuals. Piombino et al.^25^ demonstrated that saliva samples from obese individuals could more significantly suppress the release of wine aroma compared with those from the normal-weight. The authors suggested that this was likely due to the difference in protein contents. Muñoz-González et al.^26^ reported in their *ex vivo* study that saliva samples from three healthy individuals exhibited different extents of aroma retention effects, which was related to the distinct total protein content and total antioxidant capacity of their saliva samples. Sensory studies have revealed differences in wine preference amongst consumers with various cultural backgrounds. Sáenz-Navajas et al.^27^ found that the intensity of astringency was negatively correlated with the quality level given by the Spanish consumers while not with French consumers. However, we do not know if the difference in salivary composition amongst wine tasters with different cultural backgrounds can explain variations in wine perception and preference.

Wine is a high-valued agricultural product appreciated by consumers around the world. A better understanding of the preference for wine by consumers from different countries and regions could enable wine producers to promote attractive products to particular consumer groups. The present study is based on 13 Western and 13 Chinese experienced wine tasters. By performing Pivot^®^ Profile and continuous scale assessment, differences in wine perception and preference between the two groups were confirmed. Headspace solid-phase microextraction-comprehensive two-dimensional gas chromatography−mass spectrometry (HS-SPME-GC×GC−MS) analysis was carried out to identify the effect of saliva on wine aroma release. Protein compositions of stimulated saliva samples were analysed by quantitative liquid chromatography with tandem mass spectrometry (LC−MS/MS). The Western group had significantly higher proline-rich protein (PRPp), while the Chinese group had saliva more abundant in lipocalin-1 (LCN-1p). By integrating these results, we can begin to explain cultural differences in wine perception and preference by salivary protein composition.

## RESULTS AND DISCUSSION

### Experimental design

To minimise the impact of wine education which could significantly impact their expression of wine perception during the wine tasting,^4^ all participants were experienced wine tasters who were able to use English terms to describe wine characteristics. Two sensory evaluation sections captured their perceptions of wine flavour and preference for wine, which were then explained through enzyme assays and volatile and proteomics analyses. To mimic the wine tasting condition as closely as possible, significantly shorter headspace equilibrium and extraction times were adopted. To tackle the complex nature of wine, the highly sensitive and improved separation capacity of the GC×GC−MS technique was therefore used for the analysis. Comprehensive quantitative profiling of salivary proteins was done by TMT-based quantitative LC−MS/MS analysis, which provided valuable information on the identities and relative concentrations of proteins with sensory impacts during wine tasting. The integration of these data highlighted the possibility of explaining the cultural difference in wine preference by salivary protein composition.

### Sensory perception and preference of wine

Shown in **Table 2**, eight wine samples varied in basic chemical parameters. It was observed that wines from the same regions had relatively similar values, for example, wines 3 and 7 from Strathbogie ranges, and wines 5 and 6 from Pyrenees. Sugar accumulation in grapes is susceptible to temperature, which then affects the alcohol content in the resulting wine.^28^ The accumulation of tannin compounds in grapes is mostly affected by sunlight exposure.^29^ During maceration, tannin compounds originating from grape seedcoats and skins enter the wine.^30^ Since wine samples were products of different wineries, different winemaking practices will have been adopted during production. These results might suggest that the wine region’s environment is a driving factor for alcohol and tannin content in wine. In agreement, Tu et al.^31^ recently reported that tannin profiles of Cabernet Sauvignon grapes could be used to distinguish wine regions.

The Chinese group perceived significantly stronger “fruity”, “floral” or “sweet” notes from wines 1, 3, 6, 7 and 8, while the difference in the perceived intensities of investigated attributes from wines 2, 4 and 5 were not statistically significant (**Table 2**). This result might imply that the Chinese group were more sensitive to the presence of these three attributes in wine. Although the two groups did not display a significant difference in their overall liking towards most of the wines, the Chinese group showed a particular preference for wine 8, which had the highest alcohol, residual sugar and condensed tannin contents.

To better understand the differences in wine sensory perception and preference between the two groups, data obtained from the continuous scale assessment were analysed by PCA as illustrated in **Fig. 1 (a and c)**. For the Western panellists (**Fig. 1a**), 69.8% of the total variance is explained by the first two dimensions. Wine 4 is positioned in the space defined by positive Dim 1 and negative Dim 2, which is characterised by attributes including “body”, “astringent”, “umami” and “bitter”. The opposite space localised wine 1, 6, and 7. Wine 8 is found in the space defined by negative Dim 1 and negative Dim 2, which is associated with “woody” and “sweet”. However, wine samples are not distributed in the opposite space associated with “floral”, “fruity” and the degree of overall liking (label: Taste. liking). For the Chinese group (**Fig. 1c**), most of the total variance (78.8%) is explained in the first two PCA dimensions. Wine 2, 5 and 7 are predominantly separated in Dim 1 with “bitter” and “floral” as the driving attributes. Projections of the other five wine samples are mostly influenced by Dim 2. Wine 4, 6 and 8 are positioned on the negative side of the Dim 2 while wine 1 and 3 are on the positive side, mainly contributed by attributes such as “umami”, “earthy”, “spicy”, “sweet”, “body”, as well as the overall liking.

**Fig. 1.**
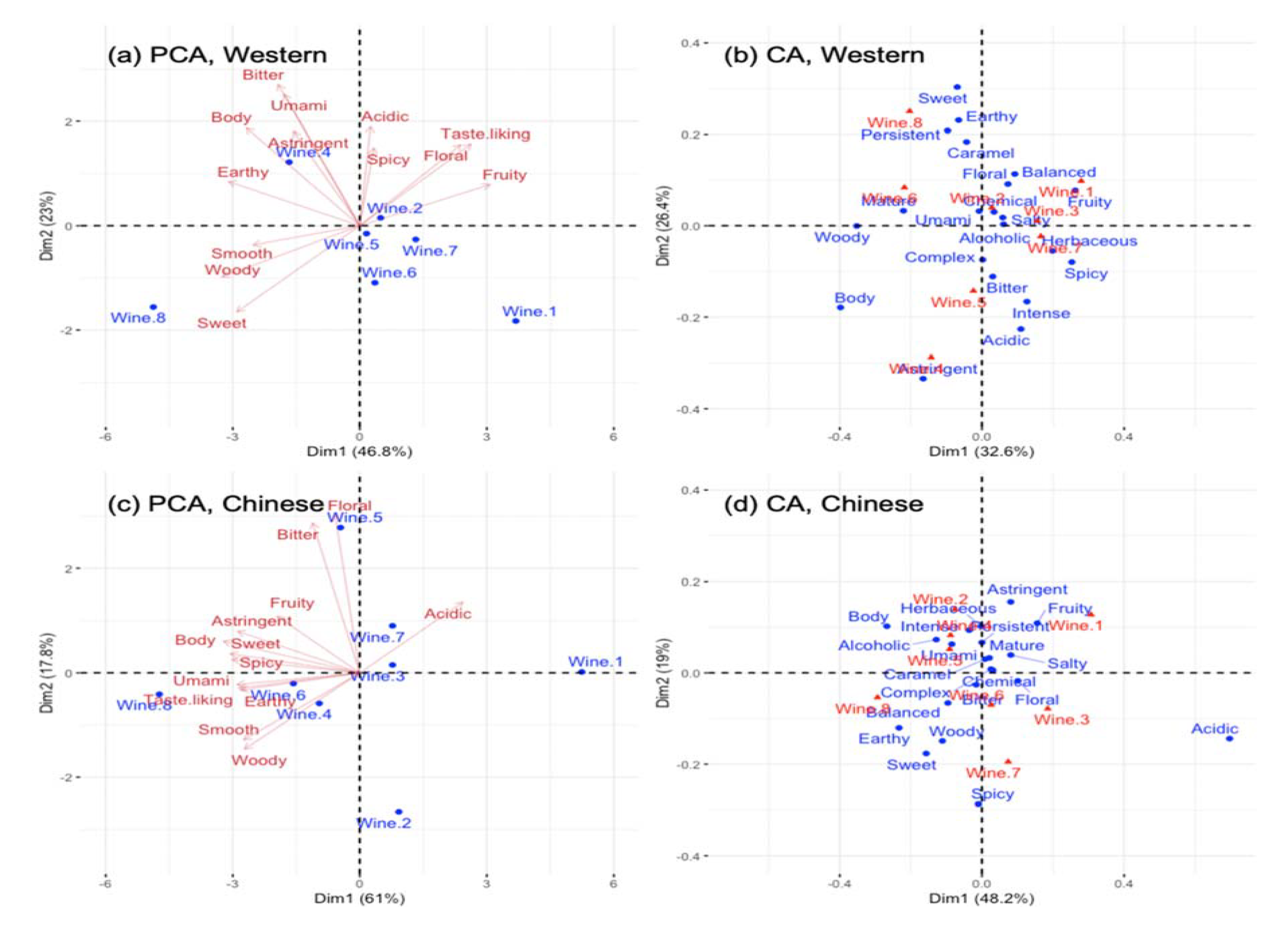
Principal component analysis (PCA) illustrating continuous scale assessment results of **(a)** Western, **c)** Chinese wine consumers and correspondence analysis (CA) illustrating translated frequency of wine iptors used by **(b)** Western, and **(d)** Chinese wine consumers.

The two groups of panellists had different wine preferences. Comparing the distribution of attributes and wine samples and that of overall liking, the Western group preferred wine with stronger “floral” notes while “sweet”, “woody” and “smooth” were not considered desirable. Yet the Chinese group preferred earthy and umami wines with low acidity. Interestingly, it was observed that wine 8 matched the Chinese group’s preference while the overall characteristics of wine 8 did not attract the Western group. These findings were consistent with the finding obtained from comparing group means of overall liking scores.

Disregarding cultural and ethnic backgrounds, wine preference is mainly determined by the sensational perception during wine tasting. The continuous scale assessment only allows panellists to rate attributes provided in the questionnaire,^32^ which may reflect the whole flavour profile perceived by the taster. Therefore, a second sensory session aimed at understanding wine perception was performed.

The result of the CA based on the contingency tables of Pivot^©^ profile revealed the variations in perception. For the Western group (**Fig. 1b**), the first two CA dimensions accounted for 59.0% of the total variance. For Dim 1, “sweet”, “earthy” and “persistent” are major descriptors contributing to the positive side while “astringent” and “acidic” contribute to the negative side. In addition, a clearer separation is observed from Dim 2, especially for wine 8 and 4, which are placed towards the positive and negative ends, respectively. Dim 2 is positively characterised by descriptors such as “Spicy”, “fruity” and “balanced” while mainly negatively characterised by “woody”. The first two DA dimensions explain 67.2% of the total variance for the Chinese group (**Fig. 1d**). In Dim 1, wine 2 and 7 are separated toward opposite ends strongly driven by “astringent” and “spicy”. The space defined by positive Dim 1 and positive Dim 2 is characterised by “fruity” localised wine 1. The space defined by negative Dim 1 and negative Dim 2 is characterised by “earthy”, “sweet” and “woody” localised wine 8. However, other sensory descriptors and wine samples were distributed around the origin.

As a free-description-based method, Pivot^©^ profile is advantageous in capturing perception during wine tasting compared to a scale assessment.^33^ Comparing the CA plots, it could be found that the two groups of tasters had some common perception to a certain extent. For example, wine 1 was described as “fruity”; wine 7 was described as “spicy” and wine 8 was described as “earthy” by both groups. Yet, considerable variation in their perceptions was observed. Typically, the Western group considered “astringent” the most typical characteristic of wine 4 while the wine impressed the Chinese group with its “intense”, “acidic” and “umami” characteristics. Moreover, despite both groups describing wine 7 as spicy, “herbaceous” is another pronounced aroma perceived by the Western group but not by the Chinese group.

Differences in wine perception and preference between the Western and Chinese could be due to the variations in cultural background and wine education. Some food products commonly used for describing wine aroma do not appear in a traditional Chinese food system. Good examples are “blackcurrant”, “raspberry” and “liquorice” and while there have been attempts to develop equivalences to translate Chinese wine descriptors to Western ones, for example, “dried Chinese hawthorn” and “blackberry preserve”, many descriptors are not interchangeable.^34^ It would be challenging to take cultural and educational factors into consideration when attempting to understand wine perception and preference across cultures by their salivary composition. Therefore, eligible participants in this study were experienced wine tasters who had received formalised wine education and were able to use Western wine descriptors properly. Our screening criterion could minimise the impacts of cultural background and education.

### Effect of panellists’ saliva on the release of wine aroma

Two groups of panellists displayed a difference in the perception of “fruitiness” and “floralness” from wine samples in both sensory evaluation sessions. As the perception of wine flavour is determined by the retronasal olfaction of volatile compounds, GC×GC−MS analysis on a wine spiked with pooled saliva samples of the two groups was conducted. A total of 25 ester, 12 alcohol, 4 aldehyde, 1 acid, 3 terpene and 2 ketone compounds were identified and semi-quantified (**Fig. 2a**). The results suggested that compared with spiking in the pooled Western saliva, wine spiked with the pooled Chinese saliva had significantly (*p < 0.05*) higher concentrations of most detected volatiles including the majority of esters, alcohols and aldehydes as well as octanoic acid and styrene. On the contrary, the pooled Western saliva only resulted in a stronger release of ethyl dodecanoate, hexyl acetate, 2,3-butanediol, 2-ethyl-1- hexanol and limonene (**Fig. 2b**).

**Fig. 2.**
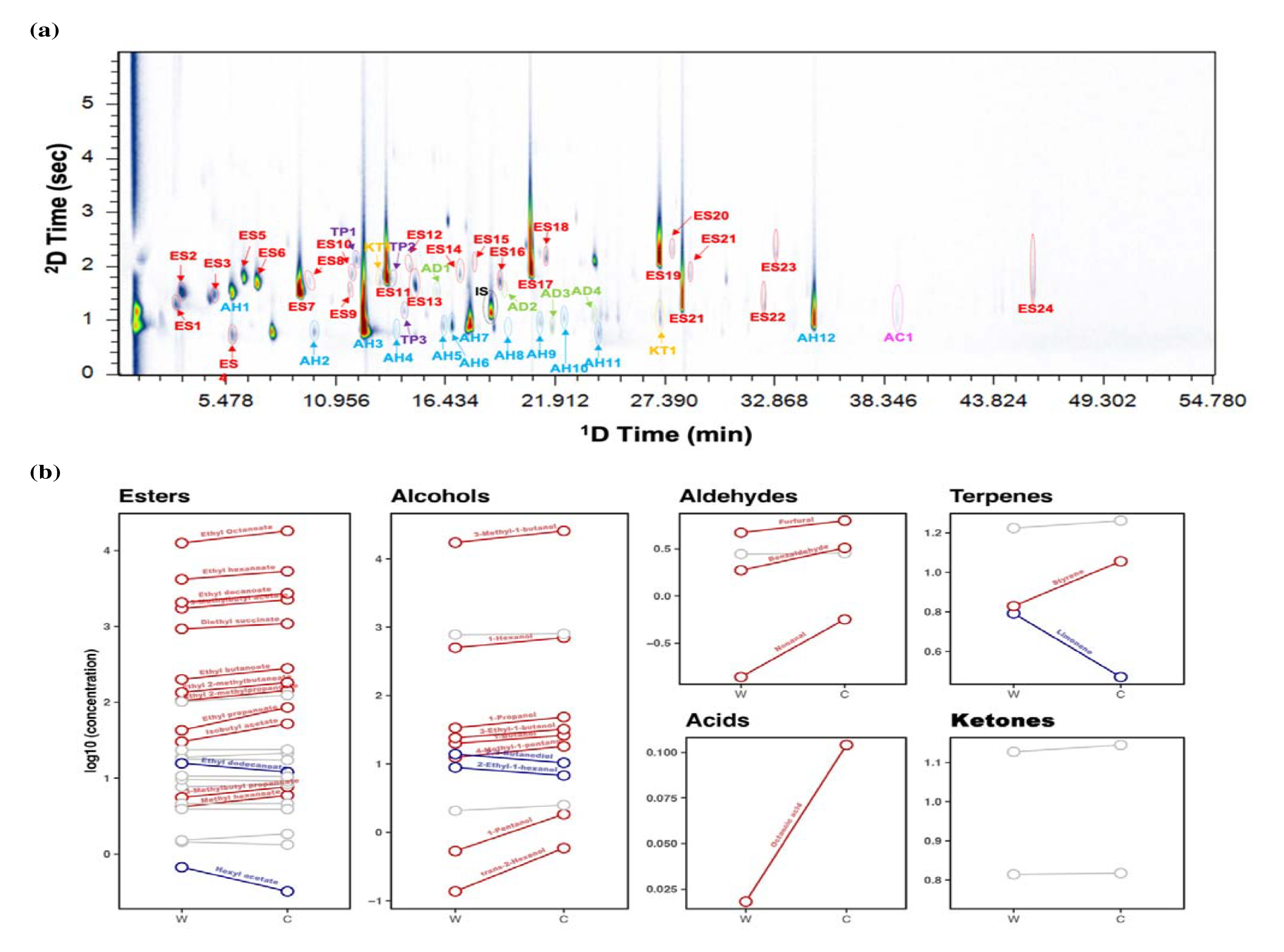
**(a)** Comprehensive two-dimensional GC data displaying compounds separated and identified in the GC−MS analysis. The internal standard (IS) is highlighted in black. Compounds are labelled according ters (ES), alcohols (AH), aldehydes (AD), acids (AC), terpenes (TP) and ketones (KT) are highlighted d, blue, green, magenta, purple and orange, respectively. ES: 1: ethyl propanoate, 2: ethyl 2- ylpropanoate, 3: isobutyl acetate, 4: ethyl butanoate, 5: ethyl 2-methylbutanoate, 6: ethyl 3- ylbutanoate, 7: 3-methylbutyl acetate, 8: ethyl pentanoate, 9: methyl hexanoate, 10: 3-methylbutyl anoate, 11: ethyl hexanoate, 12: isoamyl butanoate, 13: hexyl acetate, 14: ethyl heptanoate, 15: isobutyl noate, 16: methyl octanoate, 17: ethyl octanoate, 18: isopentyl hexanoate, 19: ethyl decanoate, 20: 3- ylbutyl octanoate, 21: diethyl succinate, 22: ethyl 9-decenoate, 23: phenylethyl acetate, 24: ethyl canoate and 25: diethyl phthalate. AH: 1: 1-propanol, 2: 1-butanol, 3: 3-methyl-1-butanol, 4: 1-pentanol, methyl-1-pentanol, 6: 3-ethyl-1-butanol, 7: 1-hexanol, 8: *trans*-2-hexenol, 9: 1-heptanol, 10: 2-ethyl-1- nol, 11: 2,3-butanediol and 12: phenylethyl alcohol. AD: 1: octanal, 2: nonanal, 3: furfural and 4: aldehyde. AC: 1: octanoic acid. TP: 1: limonene, 2: γ-terpinene and 3: styrene. KT: 1: 4-octanone and 2: olactone. **(b)** The effects on wine volatile release after mixing with Western and Chinese pooled saliva les were compared. Blue and red lines indicate the detected compounds in the headspace are significantly *0.05*) higher in the wine-Western saliva and wine-Chinese saliva mixtures, respectively. Grey line ates the difference is not significant.

Esters are a major class of wine volatile compounds, contributing to pleasant fruity notes enjoyed by consumers, therefore esters are considered critical chemical indicators for winemakers to produce wines with more pounced fruity characteristics.^35^ Like esters, alcohols are also present at high concentrations in wine. Although the detected significant alcohols themselves are associated with providing unpleasant nuances such as malt, fusel oil or solvent,^36^ they could play a role in the perception of “fruitiness”. Cameleyre, et al. ^37^ reported that 1-butanol alone could lower the olfactory threshold of “fruitiness” while together with other wine higher alcohols, it increased the perception threshold. The authors explained that the suppression effect on “fruity” notes was because higher concentrations of higher alcohols could mask the smell of less intense compounds. This odour masking effect of higher alcohols was also observed by Fuente-Blanco, et al. ^38^ Compared to esters and alcohols, other groups of volatiles present at relatively low concentrations (**Supplementary Table 3**) and some of these compounds could contribute to wine flavour perception with their low olfactory thresholds like limonene, while some such as benzaldehyde^39^ normally do not reach their odour thresholds in wine.

Salivary enzymes can influence wine aroma release in the mouth.^40^ In order to understand the differences in the ester release and thereby the “fruitiness” perception between the two groups from an enzyme activity perspective, total salivary esterase activity assay was conducted (Table 1). In agreement, the result in this study showed a significantly stronger esterase (*p < 0.05*) activity in the pooled Chinese saliva. Higher salivary esterase activity was associated with stronger hydrolysis of ester compounds in the mouth.^41,42^ Interestingly, an overall higher esterase release was found in the Chinese group, which had a strong esterase activity to hydrolyse esters. Significant interindividual differences in the salivary esterase activity have been demonstrated by María, et al. ^42^. The effect of esterases on differentiating the ester release between the two groups and potentially their perception of “fruitiness” was limited. The Chinese pooled saliva sample also exhibited stronger (*p < 0.05*) α-amylase activity yet the enzyme was not likely to contribute to the variation in volatile release between the two groups. Salivary α-amylase can influence flavour perception by inducing starch hydrolysis.^43^ In this study, this function had a negligible impact as there was no starch in wine and panellists were asked not to eat or drink 2 h prior to the study, leaving a low probability for undigested starch to remain in the mouth.

**Table 1.** Information of the Western and Chinese panellists

|  | Western<br>(*n* = 13) | Chinese<br>(*n* = 13) |
| --- | --- | --- |
| *Gender* |  |  |
| Female | 6 | 6 |
| Male | 7 | 7 |
| *Age* |  |  |
| 18–24 | 0 | 1 |
| 25–30 | 0 | 7 |
| 31–40 | 3 | 4 |
| 41–55 | 6 | 1 |
| 56+ | 4 | 0 |
| *Occupation* |  |  |
| Wine related | 8 | 5 |
| Others | 6 | 7 |
| *Basic salivary parameters* |  |  |
| Protein concentration (g/L) | 0.76±0.16 | 0.75±0.15 |
| α-Amylase activity (unit of activity per min) | 0.072±0.006 | 0.114±0.01* |
| Esterase activity (unit of activity per min) | 0.079±0.001 | 0.081±0.001* |

Salivary protein concentration, and α-amylase and esterase activities are expressed as mean ± standard deviation. Significantly different ( $p < 0.05$ ) mean values between the two groups are marked with “\*”. Enzyme activities were measured using pooled saliva samples.

Although *ex vivo* investigations on the effects of esterase and α-amylase activities of the participants did not support our observations of volatile analysis, other salivary enzymes could influence the oral volatile release by their aroma retention function. This function of enzymes, like other non-enzyme salivary proteins, is achieved by inducing hydrophobic sites where small aroma molecules are bound through the hydrophobic interaction.^10^ Since the effect of aroma retention is directly associated with the number of hydrophobic sites on enzymes, a more comprehensive understanding of the effect of these enzymes could be revealed by the quantitative analysis.

### Can variation in wine perception and preference amongst consumers be explained by salivary protein composition?

Proteomics revealed the composition and abundance of salivary proteins, which were compared between the two groups (**Fig. 3**). We identified a total of 121 proteins, and of these, the relative abundances of 83 proteins (grey dots) were not significantly different between the two groups and these include esterases and α-amylase. Thirty five salivary proteins (red dots) were significantly higher (FDR < 0.01, s0 > 2) in the Chinese group, and amongst them, lipocalin-1 (abbreviation: LCN-1, gene name: *LCN1*) was previously reported to have a potential contribution to sensory perception.^8^ In comparison, saliva samples of the Western group had significantly more abundant salivary acidic proline-rich phosphoprotein 1 (abbreviation: PRH1, gene name: *PRH1*), basic salivary proline-rich protein 2 (abbreviation: PRB2, gene name: *PRB2*), and basic salivary proline-rich protein 3 (abbreviation: PRB3, gene name: *PRB3*) (blue dots). These proteins belong to the proline-rich proteins (PRPs), which are able to bind polyphenol, forming complexes and contributing to the sensory descriptor of ‘astringency’.^44^

**Fig. 3.**
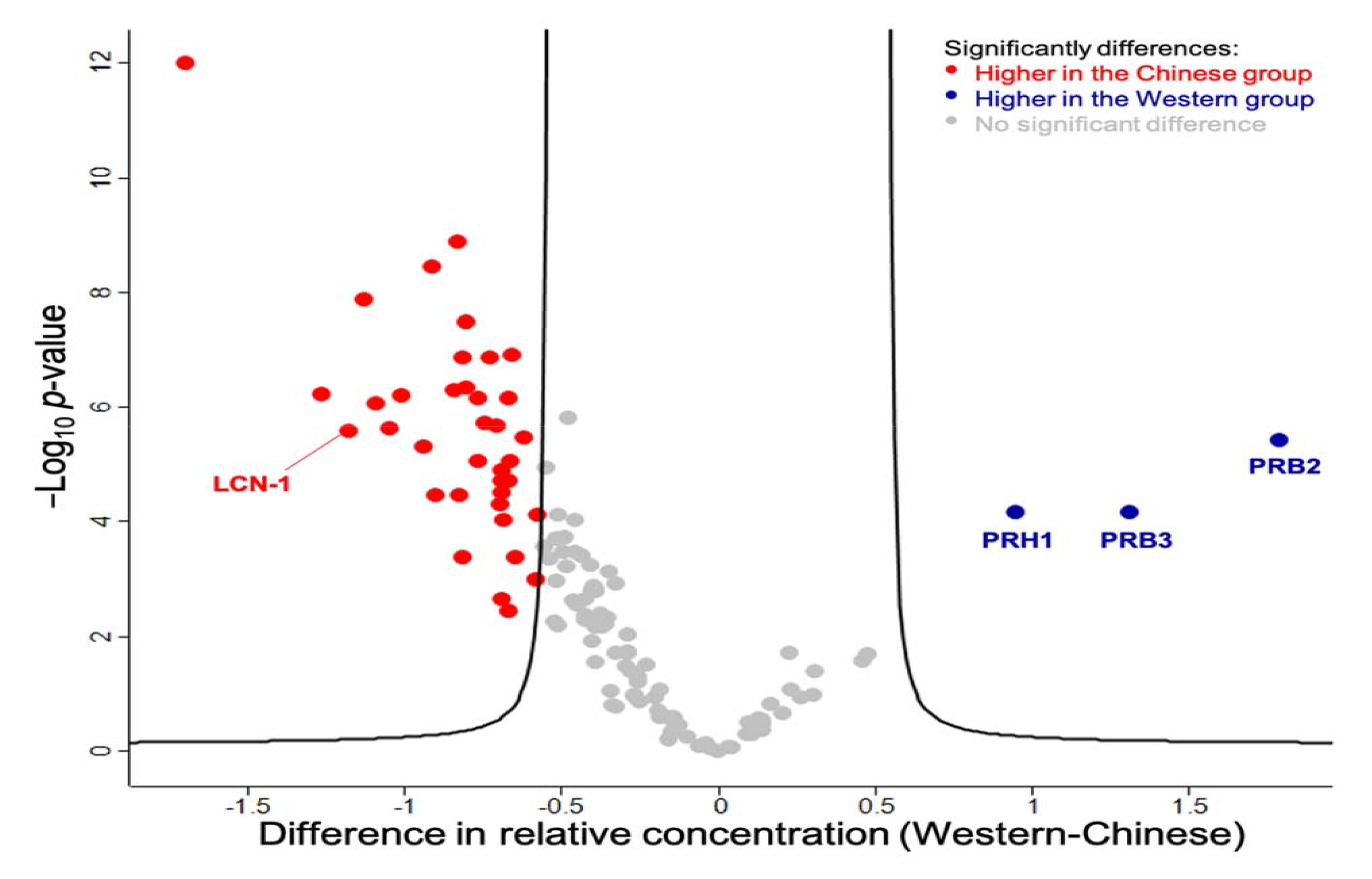
The variation in salivary protein compositions between Western and Chinese panellists. Proteins with ficant differences (FDR = 0.01, s0 = 2) between the two groups were located outside the curve. Proteins reported impacts on the sensory perception are labelled with their gene names.

Circadian rhythms, health status, exercise, oral microbiome and protease activity influence the composition of proteins in human saliva.^45^ Major variance has been found in the salivary proteome by age, gender and ethnicity.^20,46^ In this study, although salivary proteins with significantly different concentrations were discovered when comparing between genders and between different age groups, the most significant proteins were found in the comparison between ethnicities (**Supplementary Table 4**). Therefore, it is possible that variation in ethnicity was the driving reason for the difference in the saliva proteome. Mosca et al.^20^ reported that compared with the Dutch participants (*n* = 15), Chinese participants (*n* = 15) had a higher average salivary protein concentration, which could be due to the difference in diets between the two groups, yet total protein concentrations between the two groups were not significantly different in our study (Table 1). Moreover, esterases and α-amylase were not significantly different in concentrations between the two groups. Rather than the total salivary protein content, the variation in wine aroma release and potentially the perception was more likely due to the difference in the non-enzyme protein composition.

To further understand the role of these salivary proteins, selected sensory attributes which were significantly different tested in the Wilcoxon Singed Rank test, LCN-1 and three PRPs were analysed by the Pearson correlation analysis (**Fig. 4**). Results suggested that concentrations of salivary PRPs were negatively correlated with the perception of “floral” and “fruity” notes from all wine samples. Salivary PRPs exhibit a strong affinity toward tannins, which is contributed by their extended structure, making proline binding sites extremely accessible for tannins.^47^ These tannin-PRP complexes can effectively bind small molecules such as wine esters and terpenes via hydrophobic interactions.^48^ As these small molecules could contribute to the “floral” and “fruity” notes, the observed negative correlations between PRPs and the corresponding perceived intensities could be explained. The formation of tannin-PRP complexes increases the friction in the mouth and determines the perception of “astringency”.^49^ Theoretically, concentrations of PRPs and the astringency perception should be positively correlated. However, levels of PRH-1 and PRB-2 were predominantly negatively correlated with the intensity of “astringent” sensations in all wines. Other studies have showed that the correlation between the perceived intensity of astringency and PRP concentrations as ambiguous.^50-52^ The astringency sensation is complicated and requires further study to correlate perception of this descriptor with PRPs.

**Fig. 4.**
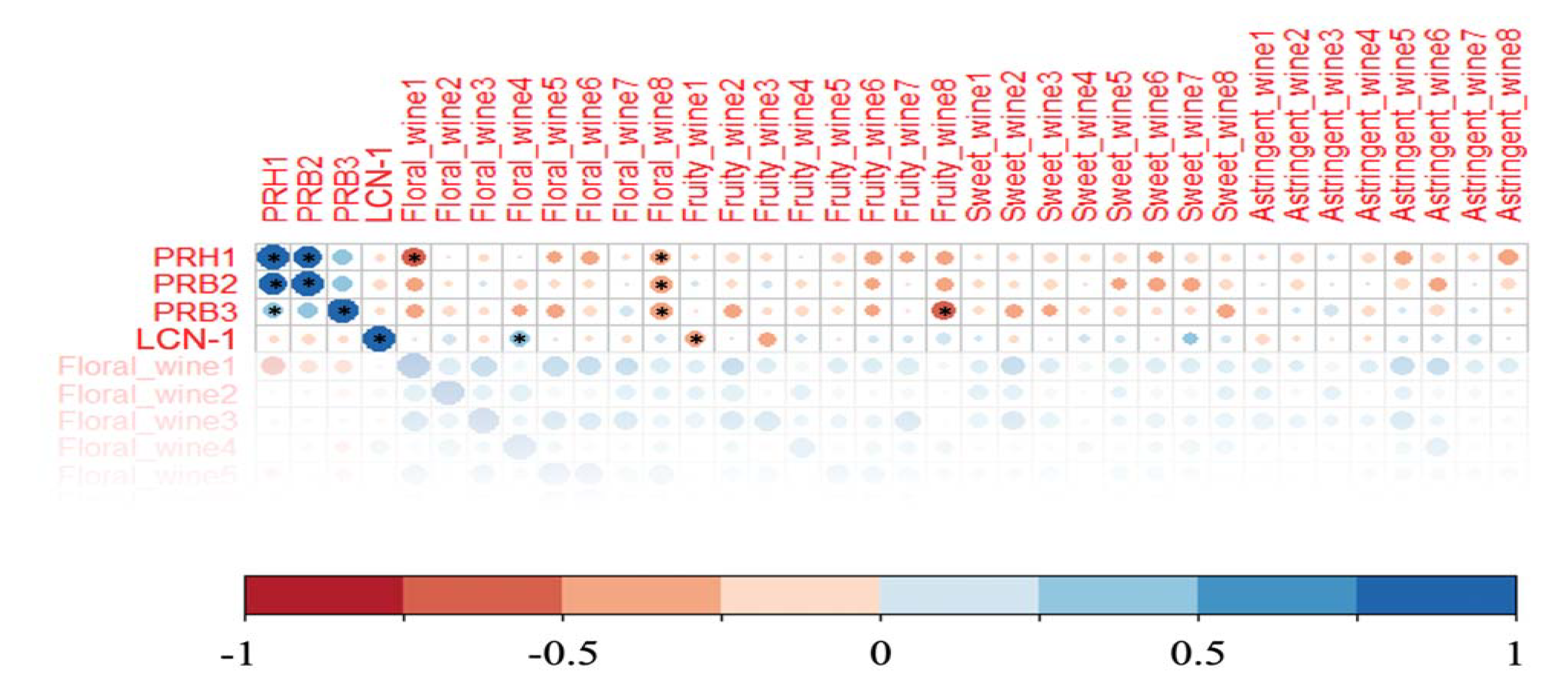
Pearson correlation analysis amongst intensities of sensory attributes (“floral”, “fruity”, “sweet” and ngent”) perceived from wine samples by panellists and relative concentrations of their salivary PRH1, 2, PRB3 and LCN-1. Significant correlations at the 0.05 level are labelled with “*”.

Lipocalin protein could potentially enhance the perception of wine aroma. This was witnessed in many positive correlations between LCN-1 concentration and perceived intensity of “fruity” and “floral” bouquets. *LCN1* encodes LCN-1 protein belonging to the lipocalin superfamily that is released by the von Ebner’s glands around the circumvallate and foliate papilla of the human tongue.^53^ In our study, LCN-1 was significantly (*p* < 0.05) negatively correlated with the perception of “fruitiness” in wine 1, perhaps explained by the difference in the matrix composition of the sample wines along with the positive correlations found for “floralness” in wines 1, 2, 4, 5 and 8 as well as “fruitiness” in wines 4, 6, 7 and 8 (**Fig. 4**). Lipocalin in the mucus of the nasal cavity is likely to be directly involved in aroma perception, but it remains unclear whether volatiles carried by salivary LCN-1 can be transported to the olfactory receptors in the nasal cavity. Ployon, et al. ^8^ emphasised another mechanism by which LCN-1 could play a role in the aroma release. Non-enzyme salivary proteins can directly influence volatile release by their binding aroma effect. However, LCN-1 only presents at approximately 100 ng/mL in human saliva,^54^ its capacity of influencing the overall volatile profile is questionable. We used β-lactoglobulin (BLG) as a proxy for LCN-1 which also belongs to the lipocalin family and has a similar structure to LCN-1. The result showed no significant difference in the volatile release of wine samples spiked with 100 ng/mL or 100 μg/mL BLG or only the buffer solution (**Supplementary Table 5**), suggesting that LCN-1 does not play a significant role in changing the composition of aroma compound in the mouth during wine tasting.

This present study reports the difference in salivary protein composition between Western and Chinese wine tasters and particular proteins that might play a role in varying the aroma release in the mouth or reacting with the olfactory receptor. Using a small panel, this information could provide new insights into variance of wine perception and preference. Of note that there exists differences in dietary patterns of consumers though they have the same ethnicity,^2,55^ and human salivary composition is susceptible to the diet.^21^

## CONCLUSION

In-mouth reactions of wine and saliva affects preference for wine. We show here that consumers with different cultural backgrounds have different salivary protein compositions. While many proteins varied between the wine tasters, the differences in abundances of salivary PRPs and LCN-1 may explain the variations in the perception of wine flavour and preference as observed in the sensory evaluation. GC×GC−MS analysis supported this observation from a volatile release perspective. We have identified a potential contribution of LCN-1 to the perception of wine taste, but we have been unable to confirm a direct mechanism by which this occurs. Our findings provide a biochemical basis to investigate flavour and aroma preferences for wine and potentially other food products by investigation of salivary proteins. Using saliva composition to achieve a better understanding of preference could enable food and beverage products to be designed and targeted to distinct consumer or population groups.

## MATERIALS AND METHODS

### Chemicals and consumables

All solvents and chemicals used were of analytical grade. Methanol, Folin-Ciocalteu reagent, concentrated hydrochloric acid sodium carbonate, gallic acid, vanillin, catechin, 1M triethylammonium bicarbonate (TEAB) buffer, sodium chloride, acetone, urea, iodoacetamide, tris(2-carboxyethyl)phosphine hydrochloride (TECP), acetonitrile (ACN), trifluoroacetic acid (TFA), β-lactoglobulin, phosphate-buffered saline, 4-octanol used in the GC analysis and amylase activity assay kit were purchased from Sigma-Aldrich (Castle Hill NSW, Australia). Micro bicinchoninic acid (BCA) protein assay kit was purchased from Thermo Fisher Scientific (Waltham, MA). Purified water was obtained from a Milli-Q system (Millipore Australia, Bayswater, Victoria, Australia). Eight Shiraz wines produced in Victoria, Australia were purchased from local wine producers. Due to the same wine samples being no longer available on the market when the spiking tests were conducted, Wine 8 of vintage 2019 was used for the spiking tests.

### Conventional chemical analysis of wine

Total residual sugar content (glucose + fructose), alcohol content, total titratable acidity (TA), pH, volatile acid content, and malic acid content were measured with an OenoFoss wine analyser (Foss Analytical, Hillerod, Denmark).

### Determination of wine total phenolic content (TPC)

Quantification of TPC in wine samples was based on the method described by Iland^56^ with some modifications. Briefly, 2 mL wine sample was mixed with 8 mL methanol to prepare a 1:5 dilution and then 50 μL of the diluted sample was mixed with 1.25 mL 0.2 N Folin-Ciocalteu reagent. After incubating at room temperature for 5 min, 1 mL saturated Na_2_CO_3_ was added and mixed well with a vortex. The mixture was incubated at 37 °C for 1 h before transferring the mixture (200 μL) to a microplate for the absorbance measurement at 765 nm with a Varioskan Flash plate reader (Multiskan FC Microplate Photometer, Thermo Scientific, Waltham, MA, USA). A calibration curve of the standard compound gallic acid with a linearity range from 50 mg/L to 800 mg/L was constructed. TPC was expressed as gallic acid equivalents (mg GAE/L wine). This assay was performed in triplicates.

### Determination of total condensed tannin content (CTC) and tannins by bovine serum albumin (BSA) precipitation assay

To determine CTC, a procedure slightly modified from Broadhurst and Jones^57^ was adopted. Fifty microlitres of the diluted sample (1:5 with Milli-Q water), 3 mL of 4% vanillin in methanol (w/v) and 1.5 mL of concentrated HCl were mixed. After 15 min, the absorbance of the mixture was measured at 500 nm. BSA precipitation assay was conducted following Harbertson, et al. ^58^ without modification. For both assays, catechin was used as the standard and results were expressed as catechin equivalents (mg CA/L wine or g CA/L wine). These assays were performed in triplicates.

### Sensory analysis panellist recruitment

Experienced wine tasters were recruited in Victoria, Australia. They were either wine professionals (e.g., wine judges, winemakers and owners of wineries) or people who acquired Wine and Spirit Education Trust (WSET) level 2 award in wines or equivalent. After screening ineligible panellists, 26 participants aged from 24 to 60 years were selected. According to their self-reported ethnicities, they were classified into Western (*n* = 13) and Chinese (*n* = 13) groups, balanced by gender. The background information of the panel is summarised in **Table 1**. Sensory panel recruitment and the use of their saliva samples for analysis were approved by the Office for Research Ethics and Integrity of the University of Melbourne (Ethics ID: 1852616). All participants gave informed written consent to take part in the study.

### Saliva collection

The saliva collection procedure was conducted using the Saliva Check Buffer kit (GC Europe N.V., Leuven, Belgium) following its instructions before the sensory evaluation session. Briefly, participants were instructed not to consume food or beverages (except water) 1 h before saliva collection. A wax gum was provided to each panellist for saliva stimulation. In a five-min collection time, participants were instructed to chew the piece of wax and expectorate into the saliva collection cup at regular intervals. After collection, saliva samples were centrifuged at 15,000 ×g for 15 min at 4 °C to remove insoluble matter and then separated in aliquots and kept frozen at −80 °C until analysis.

### Sensory evaluation of the perception of wine by the Pivot^©^ profile

Sensory evaluation sessions were conducted in the sensory laboratory at the University of Melbourne with room temperature maintained at 22±2 °C. Each booth was equipped with standard white LED lights and pass-through compartment doors for delivering samples and questionnaires. The assay was conducted based on Thuillier et al.^33^ with some modifications. A pivot wine was prepared by mixing 100 mL of each of the wine sample. Wines were only opened when pouring was necessary. After pouring, 100% pure food-grade argon gas (as claimed. Winesave^®^, VIC, Australia) was applied to the wine headspace followed by tightly sealing with parafilm to minimise oxidation. Samples were presented in random order, and each wine (40 mL) was given to panellists together with a glass of pivot wine (40 mL). Panellists were asked to complete a questionnaires comprised of a “less” and a “more” column. For a perceivable attribute, if its perceived intensity was stronger in the sample than the pivot, the attribute was filled in the “more” column, and vice versa. These attributes were then classified into 21 semantic groups including “fruity”, “floral”, “earthy”, “herbaceous”, “woody”, “caramel”, “spicy”, “alcoholic”, “intense”, “complex”, “chemical”, “balanced”, “mature”, “body”, “mature”, “persistent”, “sweet”, “acidic”, “bitter”, “salty” and “umami”. The appearance of an attribute in the “more” column contributed to a positive frequency while in the “less” column contributed to a negative frequency. For each ethnicity group, the frequencies of all semantic groups were summed. Accordingly, contingency tables of wines were built (an example is presented in **Supplementary Table 1**). For each semantic group, the balance was calculated as the difference between positive and negative frequencies. To avoid minus values, each semantic group was given a translated frequency which was the sum of the balance and the smallest absolute balance value of the group (4 and 6 for the Western and Chinese groups, respectively). The translated frequencies were then used for the correspondence analysis (CA).

### Sensory evaluation of perception and preference of wine by the continuous scale assessment

Data of measurable perceived intensities of a wide range of sensory attributes and overall liking of wine were collected using the continuous scale assessment. In this session, eight wines were presented to the panellists in random order without the pivot.

### GC×GC−MS analysis on wine spiked with pooled saliva

This assay was performed in triplicate. After thawing at 4 °C overnight, an equal volume of saliva samples from 13 Western participants was combined to prepare a pooled Western saliva sample, the same as for preparing the pooled Chinese saliva. The headspace volatile extraction method was conducted according to Muñoz-González et al.^59^ with some modifications. The 20 mL GC vial was incubated at 36 °C for 20 min. After that, 5 mL of wine, 1 mL of pooled saliva and 10 μL of 100 mg/L 4-octanol as internal standard were added to the vial followed by incubation at 36 °C for 12 min. Previous studies^59-63^ suggested that there was a significant effect of different matrixes on the GC response of internal standards, therefore the compound absolute peak areas were used for quantification. However, peak areas of 4-octanol in the wine sample spiked with the two pooled saliva samples were not significantly different in our pilot trial (**Supplementary Table 2**). In our assay, the internal standard was added for more precise quantification. A 65 μL polydimethylsiloxane/divinylbenzene (PDMS/DVB) SPME fibre with 1 cm fibre length (Supelco, Bellefonte, PA) was used to perform the headspace volatile extraction in the GC vial at 36 °C for 5 min. The efficiency of the selected SPME fibre has been tested in our lab and the fibre was used in previous publications.^64,65^

**Table 2.** Background information and comparison between perceptions of two groups of panellists for the Shiraz wines used in the nsory evaluation.

|  | Vintage | Region | pH | Alcohol<br>(% v/v) | Residual<br>sugar<br>(g/L) | Total<br>phenolic<br>(mg/L<br>GAE) | Condensed<br>tannin<br>(g/L CE) | BSA<br>tannin<br>(mg/L<br>CE) | Sensory perception between Chinese and Western<br>panellists |
| --- | --- | --- | --- | --- | --- | --- | --- | --- | --- |
| Wine 1 | 2017 | Mornington<br>Peninsula | 3.6 | 13.2 | 2.7 | 1058.2 | 6.3 | 319.6 | Stronger **fruity** and **acidic** tastes were perceived in the Chinese group while a more **astringent** mouth feel was perceived in the Western group |
| Wine 2 | 2017 | Geelong | 3.8 | 13.1 | 2.9 | 923.1 | 6.1 | 205.6 | No significant difference |
| Wine 3 | 2016 | Strathbogie<br>Ranges | 3.7 | 14.5 | 3 | 1080.6 | 6.0 | 337.8 | **Sweeter** taste was perceived in the Chinese group |
| Wine 4 | 2016 | Yarra<br>Valley | 3.7 | 15.0 | 3.4 | 1076.3 | 6.4 | 356.5 | No significant difference |
| Wine 5 | 2015 | Pyrenees | 3.6 | 13.9 | 3.2 | 1446.6 | 6.7 | 682.6 | No significant difference |
| Wine 6 | 2015 | Pyrenees | 3.7 | 14.2 | 3.0 | 1310.5 | 6.6 | 539.0 | Stronger **floral** and **sweeter** taste and **smoother** mouth feel were perceived in the Chinese group |
| Wine 7 | 2016 | Strathbogie<br>Ranges | 3.8 | 14.4 | 3.3 | 1105.4 | 7.3 | 245.6 | **Sweet** taste was perceived in the Chinese group |
| Wine 8 | 2016 | Gippsland | 3.6 | 15.2 | 4 | 1265.4 | 7.4 | 517.8 | Stronger **floral** and **fruity** taste was perceived in the Chinese group and they had **higher liking** towards this wine. |

GC×GC−MS analysis was carried out with an Agilent 7890A GC system (Agilent Technologies, Mulgrave, VIC, Australia) equipped with an SSM1800 solid state modulator (J&X Technologies Co. Ltd., Nanjing, China), coupled to an Agilent 5975C MS (Agilent Technologies). The fibre was manually desorbed in the spilt mode with a 2:1 split ratio at 220 °C for 5 min. Chromatographic separation was achieved with a SUPELCOWAX 10 (30 m × 0.2 mm internal diameter (I.D.) × 0.2 μm film thickness (*d*_f_); Supelco) as the first dimension (^1^D) column and VF-17ms (1 m × 0.1 mm I.D. × 0.1 μm *d*_f_; Agilent Technologies) as the second dimension (^2^D) column with helium carrier gas at a flow rate of 1.0 mL/min (99.999% purity). After holding at 40 °C for 4 min, the GC oven temperature was ramped up to 220 °C at a rate of 3 °C/min. The temperature was then held at 220 °C for 10 min. For the modulator, entry and exit temperatures were set the same as the GC oven program. The trap was started at −40 °C and held for 10 min. After that, the temperature was ramped to 20 °C at 1.5 °C/min and held for 5 min. The modulation period was 6 s. For MS, electron ionisation (EI) mode at 70 eV was used to obtain mass spectra with a scanning mass range from *m/z* 50 to 300, acquiring at 25.9 scans/s. MS interface, source and quadrupole temperatures were 240, 230 and 150 °C, respectively.

Alkane standards (C_6_−C_30_) were analysed to calculate retention indices (RIs) on the ^1^D column. Compound identification was facilitated with NIST library version 17 and the ^1^D RIs of compounds. Visualisation of the 2D chromatogram was achieved using Canvas version 1.5.14 (J&X Technologies Co. Ltd.). Peaks were semi-quantified facilitated with the internal standard, and concentrations were expressed as μg/L 4-octanol equivalence. Since the peak of a compound was modulated into several modulated peaks, the peak area of the compound was calculated by summation of the peak areas of the modulated peaks. Details of compound identification and quantification are provided in **Supplementary Table 3**.

### Analysis of enzyme activities on saliva

Due to the limited amount of saliva samples collected, only pooled saliva samples were subject to the enzyme activity assays. α-Amylase activity assay was carried out using an amylase activity assay kit following the user manual (Sigma-Aldrich). Total salivary esterase activity was tested following María, et al. ^42^ without modification. Assays were performed in triplicate and results were expressed as units of enzyme activity per min.

### Identification and quantitation of salivary proteins by Tandem Mass Tag (TMT) based quantitative proteomics

After removing insoluble matter in the saliva by centrifugation, 120 μL supernatant was diluted with 120 μL 50 mM TEAB in 300 mM NaCl containing protease inhibitor cocktail (Merck & Co., New Jersey, USA). Proteins in 200 μL of the mixture were participated by 1 mL pre-chilled acetone at −20 °C overnight. After centrifugation, the crude protein pellet was dissolved in 100 μL 8 M urea in 50 mM TEAB. For all samples, protein concentration was standardised to 20 μg/mL according to the protein quantification results tested by the Micro BCA protein assay kit (Thermo Fisher Scientific). Reduction was carried out using 10 μL TCEP was added followed by incubation at 37 °C for 45 min followed by alkylation with 55 mM iodoacetamide. The sample was diluted to 1 M urea with 25 mM TEAB before digestion. Sequence grade modified trypsin (1:50, enzyme:substrate, w/w) was then added. The enzymatic digestion was conducted at 37 °C overnight with agitation. After deactivating the enzyme with pure formic acid, the solution was ready for solid-phase extraction (SPE) clean-up using Oasis HLB Cartridge (Waters Corporation, Milford, MA). The cartridge was activated by passing through 1 mL 80% acetonitrile containing 0.1% trifluoroacetic acid (TFA) followed by 1.2 mL 0.1% TFA twice. After that, the sample solution was loaded and then eluted twice with 800 μL 80% ACN containing 0.1% TFA. The two extract fractions were combined and were concentrated by the speedy-vac concentrator for 20 min and then freeze-dried overnight.

Tandem mass tag (TMT) labelling was performed using the TMT10plexTM Mass Tag Labelling Kit (Thermo Fisher Scientific) with minor modifications from the manufacturer’s protocol. Individual samples were labelled in quadruplicates. Prior to labelling, a pooled sample was prepared by mixing an equal volume of samples resuspended in 100 mM TEAB and designated as the labelled pool channel (channel 126) and used as an internal control. Four hundred microlitres of anhydrous ACN were added to vials containing different isobaric labelling reagents. In new Eppendorf tubes, 10 μL of the resuspended samples, including the pooled sample, and 7 μL of the TMT labelling solution were combined. The mixture was incubated at room temperature for 1 h followed by adding 5 μL 0.5% hydroxylamine to terminate the reaction. Multiplexing of sample were then carried out by mixing 10 μL of 9 labelled samples and a labelled pooled sample. All combined samples were then freeze-dried overnight. After resuspending the freeze-dried samples with 100 μL 2% ACN containing 0.05% TFA, before LC−MS/MS analysis.

### LC-MS/MS analysis

Samples were analysed by LC-MS/MS using Orbitrap Lumos mass spectrometer (Thermo Scientific) fitted with nanoflow reversed-phase-HPLC (Ultimate 3000 RSLC, Dionex). The nano-LC system was equipped with an Acclaim Pepmap nano-trap column (Dionex – C18, 100 Å, 75 μm × 2 cm) and an Acclaim Pepmap RSLC analytical column (Dionex – C18, 100 Å, 75 μm × 50 cm). Typically for each LC-MS/MS experiment, 1 μL of the peptide mix was loaded onto the enrichment (trap) column at an isocratic flow of 5 μL/min of 3% CH3CN containing 0.1% formic acid for 6 min before the enrichment column is switched in-line with the analytical column. The eluents used for the LC were 5% DMSO/0.1% v/v formic acid (solvent A) and 100% CH3CN/5% DMSO/0.1% formic acid v/v. The gradient used was 3% B to 20% B for 95 min, 20% B to 40% B in 10 min, 40% B to 80% B in 5 min and maintained at 80% B for the final 5 min before equilibration for 10 min at 3% B prior to the next analysis.

The mass spectrometer was operated in positive-ionization mode with spray voltage set at 1.9 kV and source temperature at 275°C. Lockmass of 401.92272 from DMSO was used. The mass spectrometer was operated in the data-dependent acquisition mode MS spectra scanning from m/z 350-1550 at 120000 resolution with AGC target of 5e5. The “top speed” acquisition method mode (3 sec cycle time) on the most intense precursor was used whereby peptide ions with charge states ≥2-5 were isolated with isolation window of 0.7 m/z and fragmented with high energy collision (HCD) mode with stepped collision energy of 38 ±5%. Fragment ion spectra were acquired in Orbitrap at 15000 resolution. Dynamic exclusion was activated for 30s.

### Proteomics database search and statistical analysis

Raw data from the proteomics analysis were analysed using MaxQuant version 1.6.10.3 as described by Cox and Mann ^66^. The results were searched against the Uniprot human database (42,434 entries, June 2019) using default settings for a TMT10plex experiment with the following modifications: deamidation (NQ), oxidation of methionine and N-terminal acetylation were specified as variable modifications. Trypsin/P cleavage specificity (cleaves after lysine or arginine, even when proline is present) was used with a maximum of 2 missed cleavages. Carbamidomethylation of cysteine was set as a fixed modification. A search tolerance of 4.5 ppm was used for MS1 and 20 ppm for MS2 matching. False discovery rates (FDR) were determined through the target-decoy approach set to 1% for both peptides and proteins.

Output from the library search in the proteinGroup.txt format was processed using Perseus (version 1.6.10.0). In brief, log2-transformed TMT reporter intensities corrected by the labelled pool cannel were grouped into “Western” and “Chinese”. Filter for identification was applied to include only 100% valid values. Values were then subject to a two-sided t-test with permutation-based FDR statistics (FDR=0.01, S0=2) and the result was presented in a volcano plot following the instruction by Tyanova, et al. ^67^.

### GC×GC−MS analysis on wine spiked with β-lactoglobulin (BLG)

Phosphate-buffered saline (PBS, pH 6.7) was used to dissolve BLC at two concentrations. One millilitre of either 100 μg/mL BLG, 100 ng/mL BLG or PBS buffer was added to 20 mL GC vials containing 5 mL of wine and 10 μL of 100 mg/L 4-octanol. Incubation, HS-SPME and GC×GC−MS conditions as well as compound identification and quantification methods were identical as described in Section 2.9.

### General statistical analysis

Data analysis was performed using XLSTAT software (version 2022.2.1, Addinsoft, NY). Student’s t-test compared the means of protein concentrations and enzyme activities of the two groups. Comparisons of perceived intensities were based on Wilcoxon Singed Rank test as the sensory data were not normally distributed. Principal component analysis (PCA) and CA plots were visualised using R (version 4.2.0). Student t-test and one-way ANOVA was performed to test significant (*p* < 0.05) differences in volatile concentrations in the pooled saliva sample spiking test and the BLG spiking test, respectively.

## DATA AVAILABILITY

Datasets of this study and supplementary information are available in the repository at https://github.com/JiaqiangLuo91/datasets-and-supplementary-information.git.

## CODE AVAILABILITY

Codes used for generating the PCA and CA plots are available at: http://www.sthda.com.

## Abbreviations

HS-SPME/GC×GC−MS: headspace solid-phase microextraction/comprehensive two-dimensional gas chromatography−mass spectrometry
LC−MS/MS: liquid chromatography−tandem mass spectrometry
SPE: solid phase extraction
TPC: total phenolic content
CTC: total condensed tannin content
TA: titratable acidity
PRP: proline-rich protein
LCN-1: lipocalin-1
BLG: β-lactoglobulin protein
PRH1: salivary acidic proline-rich phosphoprotein 1
PRB2: basic salivary proline-rich protein 2
PRB3: basic salivary proline-rich protein 3
OBP: odourant-binding protein
OR: olfactory receptors
PCA: principal component analysis
CA: correspondence analysis

## ACKNOWLEDGEMENTS

The authors acknowledge the School of Agriculture and Food Sciences, Faculty of Science, the University of Melbourne for providing financial support to this study. The authors would like to thank the Melbourne Mass Spectrometry and Proteomics Facility at the Bio21 Molecular Science and Biotechnology Institute at the University of Melbourne. The authors gratefully acknowledge all experienced wine tasters who participated in the sensory evaluation sessions and donated their saliva samples. J.L. and X.R. sincerely acknowledge a Melbourne Research Scholarship offered by the University of Melbourne.

## AUTHOR CONTRIBUTIONS

Conceptualisation: J. L., P. J. M., P. Z. and K. H.; Methodology: J. L., CS. A., Y. N., P. J. M., P. Z. and K. H.; Investigation: J. L., X. R.; Data analysis and interpretation: J. L., X. R., CS. A.; Writing-original draft: J. L.; Writing-review & editing: J. L., CS. A., Y. N., P. J. M., P. Z. and K. H.; Funding acquisition: K. H.. All authors have read and agreed to the published version of the manuscript.

## ETHICAL APPROVAL

This study was approved by the Office for Research Ethics and Integrity of the University of Melbourne (Ethics ID: 1852616).

