## Supplementary files for "Variation of wine preference amongst consumers is influenced by the composition of salivary proteins"

### Supplementary Information

**Supplementary Table 1.** Contingency table of wine 1 for the Western group

| | Positive frequency | Negative frequency | Balance | Translated frequency |
|---|---|---|---|---|
| Acidic | 2 | 3 | −1 | 3 |
| Alcoholic | 1 | 0 | 1 | 5 |
| Astringent | 3 | 3 | 0 | 4 |
| Balanced | 2 | 2 | 0 | 4 |
| Bitter | 1 | 1 | 0 | 4 |
| Body | 1 | 4 | −3 | 1 |
| Caramel | 0 | 0 | 0 | 4 |
| Chemical | 0 | 0 | 0 | 4 |
| Complex | 1 | 1 | 0 | 4 |
| Earthy | 1 | 1 | 0 | 4 |
| Floral | 1 | 0 | 1 | 5 |
| Fruity | 6 | 1 | 5 | 9 |
| Herbaceous | 1 | 0 | 1 | 5 |
| Intense | 1 | 2 | −1 | 3 |
| Mature | 0 | 1 | −1 | 3 |
| Persistent | 2 | 2 | 0 | 4 |
| Salty | 0 | 0 | 0 | 4 |
| Spicy | 3 | 1 | 2 | 6 |
| Sweet | 1 | 2 | −1 | 3 |
| Umami | 0 | 0 | 0 | 4 |
| Woody | 1 | 4 | −3 | 1 |

For each semantic group, the balance was calculated as the difference between positive and negative frequencies. To avoid minus values, each semantic group was given a translated frequency which was the sum of the balance and 4 which is the smallest absolute balance value of the group (not shown in this wine sample).

**Supplementary Table 2.** Comparison between peak areas of 4-octanal in wine sample spiked with pooled Western and Chinese saliva sample.

| | Pool Western Saliva + Wine | Pooled Chinese Saliva + Wine | *p* value |
|---|---|---|---|
| Peak area | 305778 ± 45767 | 270388 ± 68696 | 0.499 |

Note: Peak areas were expressed as mean ± standard deviation. The *p* value was based on the result of Student t-test.

**Supplementary Table 3.** Comparison of the headspace relative concentrations wine volatiles after spiking for pooled Western and Chinese saliva samples.

| Compound Name | Quantifier ion | Qualifier ions | $^{1}t_R$ (min) | $^{2}t_R$ (s) | RI (exp.) | RI (NIST) | Western Saliva + Wine | Chinese Saliva + Wine |
|---|---|---|---|---|---|---|---|---|
| *Esters* | | | | | | | | |
| Ethyl propanoate | 57 | 75, 102 | 3.07 | 1.11 | 960 | 960 | 42.72±6.93 | 85.02±7.64* |
| Ethyl 2-methylpropanoate | 71 | 116, 88 | 3.37 | 1.33 | 968 | 969 | 103.84±12.27 | 142.66±17.4* |
| Isobutyl acetate | 56 | 73, 86 | 4.97 | 1.31 | 1010 | 1016 | 30.21±4.33 | 51.95±6.49* |
| Ethyl butanoate | 71 | 88, 101, 116 | 5.77 | 0.64 | 1032 | 1039 | 199.15±25.85 | 278.99±44.03* |
| Ethyl 2-methylbutanoate | 57 | 102, 85, 74 | 6.37 | 1.62 | 1047 | 1055 | 134.27±12.3 | 181.13±25.92* |
| Ethyl 3-methylbutanoate | 88 | 57, 115 | 7.07 | 1.50 | 1066 | 1072 | 101.35±11.41 | 123.7±15.63 |
| 3-Methylbutyl acetate | 70 | 55, 87 | 9.17 | 1.40 | 1120 | 1126 | 1722.27±206.02 | 2255.47±357.24* |
| Ethyl pentanoate | 85 | 57, 101 | 9.67 | 1.64 | 1133 | 1138 | 1.45±0.16 | 1.33±0.14 |
| Methyl hexanoate | 74 | 87, 59, 99 | 11.67 | 1.38 | 1185 | 1190 | 4.2±0.71 | 5.92±0.86* |
| 3-Methylbutyl propanoate | 57 | 70, 87 | 11.77 | 1.66 | 1187 | 1192 | 5.56±0.93 | 7.64±0.83* |
| Ethyl hexanoate | 88 | 99, 60 | 13.47 | 1.71 | 1232 | 1235 | 4172.99±365.54 | 5324.17±717.83* |
| Isoamyl butanoate | 71 | 55, 89 | 14.67 | 1.81 | 1263 | 1266 | 4.6±0.67 | 4.62±0.57 |
| Hexyl acetate | 56 | 61, 84, 69 | 14.97 | 1.43 | 1271 | 1274 | 0.67±0.1* | 0.32±0.04 |
| Ethyl heptanoate | 88 | 113, 101 | 17.17 | 1.67 | 1332 | 1334 | 7.66±0.59 | 8.33±1.03 |
| Isobutyl hexanoate | 99 | 56, 71, 117 | 17.87 | 1.90 | 1353 | 1351 | 1.52±0.25 | 1.84±0.27 |
| Methyl octanoate | 74 | 87, 127, 115 | 19.17 | 1.55 | 1391 | 1387 | 18.75±2.27 | 21.3±2.61 |
| Ethyl octanoate | 88 | 101, 127, 115 | 20.67 | 1.91 | 1436 | 1436 | 12544.28±1067.92 | 18173.2±2704.06* |
| Isopentyl hexanoate | 70 | 99, 55, 117 | 21.47 | 2.01 | 1461 | 1453 | 23.41±2.97 | 23.74±1.91 |
| Ethyl decanoate | 88 | 101, 155 | 27.17 | 2.08 | 1643 | 1643 | 2070.48±177.98 | 2717.55±344.18* |
| 3-Methylbutyl octanoate | 70 | 127, 145, 171 | 27.77 | 2.17 | 1663 | 1652 | 17.9±2.02 | 17.19±2.16 |
| Diethyl succinate | 101 | 129, 55, 73 | 28.27 | 1.19 | 1680 | 1681 | 926.48±63.5 | 1088.56±91.49* |

| | | | | | | | | |
|---|---|---|---|---|---|---|---|---|
| Ethyl 9-decenoate | 88 | 55, 69, 110 | 28.67 | 1.70 | 1693 | 1691 | 3.93±0.5 | 3.91±0.37 |
| Phenylethyl acetate | 104 | 91, 65 | 32.37 | 1.13 | 1826 | 1823 | 9.61±0.69 | 9.12±0.88 |
| Ethyl dodecanoate | 88 | 101, 183 | 32.97 | 2.09 | 1848 | 1848 | 15.65±1.62* | 12.03±1.05 |
| Diethyl phthalate | 149 | 177, 105 | 45.77 | 1.39 | 2373 | 2372 | 10.68±1.77 | 10.44±1.43 |
| **Total ester** | | | | | | | 22173.63±1914.6 | 30550.13±4081.07* |
| *Alcohols* | | | | | | | | |
| 1-Propanol | 59 | – | 5.77 | 1.34 | 1031 | 1037 | 33.66±5.38 | 48.24±2.14* |
| 1-Butanol | 56 | 73 | 9.97 | 0.58 | 1141 | 1144 | 19.88±1.7 | 26.21±3.95* |
| 3-Methyl-1-butanol | 55 | 70 | 12.47 | 0.75 | 1205 | 1204 | 17290.83±1619.55 | 25747.16±3395.33* |
| 1-Pentanol | 55 | 70,87 | 13.97 | 0.66 | 1245 | 1246 | 0.53±0.1 | 1.83±0.19* |
| 4-Methyl-1-pentanol | 55 | 69, 84 | 15.97 | 0.70 | 1309 | 1317 | 12.46±1.96 | 18.06±2.66* |
| 3-Ethyl-1-butanol | 56 | 69, 84 | 16.77 | 0.73 | 1321 | 1325 | 24.06±2.9 | 32.18±3.32* |
| 1-Hexanol | 56 | 69, 84, 102 | 17.67 | 0.77 | 1347 | 1348 | 499.99±53.61 | 701.84±94.85* |
| *trans*-2-Hexenol | 57 | 82, 100 | 19.57 | 0.77 | 1403 | 1401 | 0.14±0.01 | 0.59±0.06* |
| 1-Heptanol | 70 | 56, 83, 98 | 21.17 | 0.84 | 1451 | 1452 | 2.07±0.36 | 2.51±0.37 |
| 2-Ethyl-1-hexanol | 57 | 70, 83, 98 | 22.37 | 0.88 | 1488 | 1489 | 8.87±0.57* | 6.76±0.93 |
| 2,3-Butanediol | 57 | 75, 90 | 24.07 | 0.61 | 1542 | 1542 | 13.89±2.35* | 10.39±0.5 |
| Phenylethyl alcohol | 91 | 122, 65, 51 | 34.87 | 0.85 | 1899 | 1902 | 779.18±63.22 | 800.64±90.11 |
| **Total alcohol** | | | | | | | 18685.54±1731.81 | 27396.42±3581.18 |
| *Aldehydes* | | | | | | | | |
| Octanal | 56 | 84, 69, 100 | 15.57 | 1.36 | 1287 | 1290 | 2.79±0.4 | 2.85±0.27 |
| Nonanal | 57 | 70, 98, 82 | 19.37 | 1.47 | 1397 | 1397 | 0.14±0.01 | 0.57±0.07* |
| Furfural | 96 | 95, 67 | 21.77 | 0.70 | 1470 | 1471 | 4.7±0.74 | 6.3±0.22* |
| Benzaldehyde | 77 | 106, 51 | 23.77 | 0.88 | 1532 | 1532 | 1.87±0.32 | 3.24±0.37* |
| **Total aldehyde** | | | | | | | 9.5±1.26 | 12.95±0.54* |
| *Acid* | | | | | | | | |
| Octanoic acid | 60 | 73, 101, 85 | 38.97 | 0.84 | 2080 | 2087 | 1.04±0.16 | 1.27±0.07* |

| | | | | | | | | |
|---|---|---|---|---|---|---|---|---|
| **Total acid** | | | | | | | 1.04±0.16 | 1.27±0.07* |
| *Terpenes* | | | | | | | | |
| Limonene | 68 | 93, 107, 121 | 12.07 | 1.92 | 1195 | 1199 | 6.2±0.89* | 2.96±0.42 |
| γ-Terpinene | 93 | 136, 121, 77 | 13.87 | 1.91 | 1242 | 1245 | 16.75±1.26 | 18.25±2.59 |
| Styrene | 104 | 78, 51, 63 | 14.37 | 0.99 | 1255 | 1256 | 6.75±0.97 | 11.37±1.46* |
| **Total terpene** | | | | | | | 29.71±2.75 | 32.58±3.3 |
| *Ketones* | | | | | | | | |
| 4-Octanone | 57 | 71, 85, 128 | 13.17 | 1.57 | 1224 | 1224 | 6.53±1.11 | 6.57±0.86 |
| Butyrolactone | 86 | 56 | 27.17 | 0.87 | 1643 | 1647 | 13.4±1.4 | 13.96±1.68 |
| **Total ketone** | | | | | | | 19.93±2.45 | 20.53±2.25 |

Abbreviations: $^{1}t_R$: First dimension retention time, $^{2}t_R$: Second dimension retention time, RI: Retention index.

Note: RI (exp.) values (Van den Dool and Kratz) were calculated based the retention times of a series of alkane standards and the retention time of each compound on the $^{1}$D column. RI (NIST) values were extracted from the database of the National Institute of Standards and Technology (NIST17). Volatile concentrations are expressed as mean ± standard deviation of three replicates in μg/L 4-octanol equivalent. Values significantly higher ($p < 0.05$) than the other group tested by Student t-test are marked with "*".

**Supplementary Table 4.** Salivary proteins with significantly (FDR = 0.01, s0 = 2) different relative concentrations in different comparisons.

| **Ethnicity** | |
|---|---|
| *Western (n = 13)* | *Chinese (n = 13)* |
| Basic salivary proline-rich protein 2 | Immunoglobulin heavy variable 3-15 |
| Basic salivary proline-rich protein 3 | Immunoglobulin J chain |
| Salivary acidic proline-rich phosphoprotein1 | Ig kappa chain C region |
| | Ig gamma-2 chain C region |
| | Ig gamma-3 chain C region |
| | Ig mu chain C region |
| | Catalase |
| | Protein S100-A8 |
| | Myeloperoxidase |
| | Protein S100-A9 |
| | Glucose-6-phosphate isomerase |
| | L-lactate dehydrogenase B chain |
| | Heat shock protein HSP 90-alpha |
| | Neutrophil elastase |
| | Immunoglobulin alpha-2 heavy chain |
| | Ig gamma-1 chain C region |
| | Immunoglobulin lambda-like polypeptide 5 |
| | Ig lambda-1 chain C regions |
| | Immunoglobulin lambda constant 3 |
| | Glucose-6-phosphate 1-dehydrogenase |
| | Plastin-2 |
| | Pyruvate kinase PKM |
| | Myeloblastin |
| | Protein S100-P |
| | Protein S100-A4 |
| | Transketolase |
| | Lipocalin-1 |
| | Coronin-1A |

| | |
|---|---|
| | Transaldolase<br>Neutrophil defensin 3<br>Actin, cytoplasmic 2<br>Lysozyme C<br>Histone H3.2<br>BPI fold-containing family B member 1<br>Deleted in malignant brain tumors 1 protein |
| **Gender** | |
| *Female (n = 12)* | *Male (n = 14)* |
| Salivary acidic proline-rich phosphoprotein 1 | Cystatin-A<br>Immunoglobulin J chain<br>Ig alpha-1 chain C region<br>Glutathione S-transferase P<br>Immunoglobulin alpha-2 heavy chain<br>Protein S100-A12<br>Calcitermin |
| **Age groups** | |
| *25−30 (n = 7; W = 0, C = 7 / F = 4, M = 3)* | *31−40 (n = 7; W = 3, C = 4 / F = 4, M = 3)* |
| – | Basic salivary proline-rich protein 2 |
| *25−30 (n = 7; W = 0, C = 7 / F = 4, M = 3)* | *41−55 (n = 7; W = 6, C = 1 / F = 3, M = 4)* |
| Lipocalin-1<br>Lysozyme C | Basic salivary proline-rich protein 2 |
| *25−30 (n = 7; W = 0, C = 7 / F = 4, M = 3)* | *56+ (n = 4; W = 4, C = 0 / F = 1, M = 3)* |
| – | – |
| *31−40 (n = 7; W = 3, C = 4 / F = 4, M = 3)* | *41−55 (n = 7; W = 6, C = 1 / F = 3, M = 4)* |
| Protein S100-A8<br>Protein S100-A9<br>Lipocalin-1<br>Lysozyme C | – |
| *31−40 (n = 7; W = 3, C = 4 / F = 4, M = 3)* | *56+ (n = 4; W = 4, C = 0 / F = 1, M = 3)* |
| Haptoglobin<br>Ig gamma-2 chain C region | – |

| | |
|---|---|
| Ig gamma-3 chain C region | |
| Apolipoprotein A-I | |
| Protein S100-A8 processed | |
| Protein S100-A9 | |
| Neutrophil elastase | |
| Leukotriene A-4 hydrolase | |
| Ig gamma-1 chain C region | |
| Glucose-6-phosphate 1-dehydrogenase | |
| Plastin-2 | |
| Myeloblastin | |
| Protein S100-A4 | |
| Transketolase | |
| Coronin-1A | |
| Rho GDP-dissociation inhibitor 2 | |
| Neutrophil defensin 3 | |
| Actin | |
| cytoplasmic 2 | |
| Protein S100-A12 | |
| Zymogen granule protein 16 homolog B | |
| *41−55 (n = 7; W = 6, C = 1 / F = 3, M = 4)* | *56+ (n = 4; W = 4, C = 0 / F = 1, M = 3)* |
| – | – |

Abbreviations: W: Western, C: Chinese, F: Female and M: Male.

**Supplementary Table 5.** Wine headspace volatile concentrations after spiking in BLG at different concentrations.

| Compound Name | Buffer + Wine | 100 µg/L BLG + Wine | 100 ng/L BLG + Wine |
|---|---|---|---|
| *Esters* | | | |
| Ethyl propanoate | 67.99±7.38 | 65.24±8.17 | 65.83±5.22 |
| Ethyl 2-methylpropanoate | 90.01±9.89 | 92.07±12.02 | 94.23±22.87 |
| Isobutyl acetate | 31.62±2.74 | 31.46±5.16 | 30.68±4.69 |
| Ethyl butanoate | 191.67±11.81 | 208.79±33.52 | 185.99±40.47 |
| Ethyl 2-methylbutanoate | 142.45±22.98 | 157.1±31.02 | 166.67±41.66 |
| Ethyl 3-methylbutanoate | 100.92±9.7 | 102.58±19.65 | 99.26±26.2 |
| 3-Methylbutyl acetate | 1544.11±141.27 | 1702.79±273.33 | 1725.38±253.88 |
| Ethyl pentanoate | 1.39±0.26 | 1.55±0.32 | 1.54±0.13 |
| Methyl hexanoate | 4.65±0.15 | 5.54±0.88 | 5.43±1.28 |
| 3-Methylbutyl propanoate | 6.9±0.59 | 7.65±0.9 | 7.93±1.55 |
| Ethyl hexanoate | 2842.28±116.12 | 2968.02±331.67 | 3145.33±334.37 |
| Isoamyl butanoate | 4.77±0.42 | 5.71±1.07 | 5.36±0.93 |
| Hexyl acetate | 1.22±0.24 | 1.09±0.27 | 1.38±0.12 |
| Ethyl heptanoate | 6.28±0.73 | 6.9±0.48 | 6.4±0.36 |
| Isobutyl hexanoate | 1.5±0.2 | 1.75±0.14 | 1.61±0.28 |
| Methyl octanoate | 16.18±1.38 | 18.44±1.58 | 16.9±3.01 |
| Ethyl octanoate | 7847.95±268.08 | 7898.38±539.57 | 8740.15±691.63 |
| Isopentyl hexanoate | 19.84±1.69 | 21.87±1.61 | 22.26±2.36 |
| Ethyl decanoate | 1843.49±149.64 | 1918.94±111.93 | 1869.49±191.12 |
| 3-Methylbutyl octanoate | 16.02±2.38 | 18.66±2.4 | 16.56±2.97 |
| Diethyl succinate | 885.72±144.23 | 942.52±71.01 | 879.84±54.31 |
| Ethyl 9-decenoate | 4.88±1.16 | 5.62±0.39 | 5.76±0.66 |
| Phenylethyl acetate | 8.67±1.25 | 9.22±0.35 | 9.05±1.13 |
| Ethyl dodecanoate | 11.96±1.13 | 14±0.84 | 12.44±1.43 |
| Diethyl phthalate | 7.84±0.97 | 8.55±1.36 | 7.64±1.15 |
| **Total ester** | 15700.34±522.46 | 16213.81±1304.65 | 17123.08±1523.78 |
| *Alcohols* | | | |
| 1-Propanol | 30±1.54 | 33.37±2.69 | 32.26±6.96 |
| 1-Butanol | 8.26±0.75 | 8.52±1.23 | 8.81±2.27 |
| 3-Methyl-1-butanol | 10623.3±1297.51 | 11375.04±1602.46 | 11873.88±1209.25 |
| 1-Pentanol | 1.88±0.33 | 1.86±0.3 | 1.92±0.24 |
| 4-Methyl-1-pentanol | 13.63±0.44 | 14.79±2.05 | 12.86±4.52 |
| 3-Ethyl-1-butanol | 27.62±4.02 | 28.95±3.1 | 28.4±1.25 |
| 1-Hexanol | 523.73±22.54 | 570.03±74.2 | 555.59±38.13 |
| trans-2-Hexenol | 0.56±0.22 | 0.61±0.25 | 0.51±0.44 |
| 1-Heptanol | 3.16±0.12 | 3.7±0.24 | 3.56±0.36 |
| 2-Ethyl-1-hexanol | 4.52±0.24 | 5.13±0.39 | 3.64±2.48 |
| 2,3-Butanediol | 23.18±2.89 | 21.62±3.08 | 19.89±3.21 |
| Phenylethyl alcohol | 796.99±91.01 | 840.39±34.91 | 768.67±102.56 |

| | | | |
|---|---|---|---|
| **Total alcohol** | 12054.96±1255.81 | 12902.14±1702.35 | 13308.07±1143.17 |
| *Aldehydes* | | | |
| Octanal | 3.29±0.25 | 3.27±0.35 | 3.35±0.49 |
| Nonanal | 1.33±0.07 | 1.2±0.15 | 1.42±0.13 |
| Furfural | 5.13±0.51 | 5.79±0.77 | 5.42±0.72 |
| Benzaldehyde | 0.49±0.06 | 0.52±0.09 | 0.64±0.05 |
| **Total aldehyde** | 9.75±0.27 | 10.26±0.83 | 10.18±1.12 |
| *Acid* | | | |
| Octanoic acid | 1.35±0.23 | 1.8±0.21 | 1.66±0.12 |
| **Total acid** | 1.35±0.23 | 1.42±0.23 | 1.4±0.16 |
| *Terpenes* | | | |
| Limonene | 2.76±0.5 | 3.2±0.59 | 2.4±1.65 |
| γ-Terpinene | 17.41±2.46 | 18.69±1.86 | 18.51±2.75 |
| Styrene | 6.2±1.2 | 7.27±1.36 | 7.16±1.63 |
| **Total terpene** | 26.37±2.05 | 29.16±3.13 | 28.08±2.77 |
| *Ketones* | | | |
| 4-Octanone | 6.17±0.72 | 5.4±0.42 | 5.23±0.72 |
| Butyrolactone | 14.63±0.48 | 16.24±2.51 | 14.6±2.69 |
| **Total ketone** | 20.8±0.92 | 21.63±2.87 | 19.83±2.46 |

Note: volatile concentrations are expressed as mean ± standard deviation of three replicates in μg/L 4-octanol equivalent. One-way ANOVA indicated no significant ($p < 0.05$) difference in concentrations amongst the tested groups
